# Mitochondrial-nuclear variation for metabolic plasticity with potential consequences for invasion success in New Zealand mud snails

**DOI:** 10.1101/2023.09.02.556044

**Authors:** Omera B. Matoo, Humu Mohammed, Himani Patel, Maurine Neiman, Kristi L. Montooth

## Abstract

Metabolic rate is an emergent organismal trait that integrates energetic costs of living and influences species distributions and response to climate change. For ectotherms, metabolic rate is likely shaped by interactions between mitochondrial and nuclear genomes and environmental temperature. Variation in reproductive mode (e.g., asexual vs. sexual reproduction) can enhance or interfere with cross-generational transmission of these interactions, with implications for evolutionary responses to selection on standing genetic variation for metabolism-related traits. We leveraged known patterns of mitochondrial-nuclear discordance in a global invader, the snail *Potamopyrgus antipodarum*, to measure thermal plasticity of metabolic rate in invasive vs. native asexual lineages that combined multiple mitochondrial haplotypes with nuclear genomic variation from distinct lake populations. Native lineages harbored significant mitochondrial-nuclear variation for metabolic plasticity, with some genotypes maintaining low metabolic rates at high temperature. Invasive lineages contained only a small subset of this variation, with non-plastic and lower metabolic rates relative to native lineages at high temperature. Together, these data indicate that mitochondrial-nuclear-environment interactions contribute to variation in the metabolic rate of asexual lineages. Our results also demonstrate that invasive lineages have metabolic phenotypes that could facilitate successful colonization of aquatic habitats via decreased maintenance metabolism under high temperatures.

## INTRODUCTION

‘Maintenance metabolism’- a measure of minimum energy required by an organism to stay alive [1] – is one of the most fundamental physiological traits. In organisms that show normal spontaneous activity, maintenance metabolism is often termed “routine metabolic rate” (RMR). RMR is an important component of the total energy budget of organisms [1, 2], accounting for up to 50% of an individual’s total daily energy expense [1–3]. Inter- and intraspecific variation in metabolic rates has been documented to respond to a variety of environmental conditions, including temperature [2, 4].

Temperature significantly affects biochemical and physiological rates in ecotherms and therefore plays a critical role in shaping RMR, which is a particular concern in the face of climate change [2, 5]. Temperature-induced changes in physiology and metabolic rates of ectotherms in warming environments can reduce energetic tolerance to heat - a phenomenon known as metabolic meltdown [6]. This phenomenon is considered a potential driver of population failure and local extinction of ecotherms under climate change [6]. Metabolic meltdown can be countered by the ability of at least some ecotherms to remodel their physiology to reduce the effects of temperature, a form of metabolic rate plasticity that could potentially confer resilience to climate change [1, 4].

In ectotherms, plasticity of metabolic rates may (initially) manifest as an acute response to temperature, but thermal acclimation can reduce the magnitude of this response during more chronic exposures during the life span of an organism via thermal compensation [4]. If thermal compensation via plasticity were perfect, individual organisms could maintain fitness across wider temperature ranges than counterparts that show limited or no plasticity [4, 7–8]. As such, greater metabolic plasticity has been frequently suggested in the context of across-species variation in resilience to climate change [9] and successful biological invasions [10]. We nevertheless still know very little about the genomic causes and fitness consequences of intraspecific variation in these key metabolic and resilience traits in nature. Quantifying and incorporating this variation in metabolism into our models is thus critical for making predictions about the ecological and evolutionary responses of organisms to climate change.

We do know that metabolic rate varies within species [1–4, 11] and that this variation is associated with variation in nuclear DNA (nDNA) and mitochondrial DNA (mtDNA), and with epistasis between mitochondrial and nuclear genomes across a diversity of plant and animal taxa [1, 12–13]. Given the importance of metabolic rate to fund maintenance, activity, and reproduction, it is not surprising that both positive and purifying selection shape molecular variation and divergence in mitochondrial and nuclear genomes [14, 15 and references therein]. Yet, we have few systems where we understand the relationship between genetic variation in metabolic rate, the environment, and fitness well enough to predict how evolution will shape metabolic rate [2].

Reproductive mode variation (e.g., sexual vs. asexual reproduction; selfing vs. outcrossing) is common both within and across many eukaryotic taxa and is expected to influence the evolutionary dynamic between mitochondrial and nuclear (co)evolution [16, 17]. Sexual reproduction is characterized by uniparental (usually maternal) transmission of the mitochondrial genome, and recombination both within the nuclear genome and between the mtDNA and nDNA breaks up mitochondrial-nuclear (mitonuclear henceforth) combinations each generation [15]. Asexual reproduction, on the other hand, is usually characterized by inheritance of complete unrecombined mitochondrial and nuclear genomes, which will maintain mitonuclear combinations (high- or low-fitness) and is predicted to reduce the efficacy of natural selection on both genomes [15]. Clonal selection among asexual lineages could nevertheless favor lineages that capture high-fitness combinations of mitonuclear variation [15].

*Potamopyrgus antipodarum* is a New Zealand freshwater snail that is well suited for addressing questions regarding reproductive mode variation effects on mitonuclear evolutionary processes because it is characterized by natural frequent transitions from the obligately sexual diploid ancestral state to a derived obligately asexual polyploid (triploid and tetraploid) state [18]. Many studies have revealed that multiple separately derived asexual lineages coexist with sexual conspecifics in New Zealand populations [e.g., 19-20]. In its native range in New Zealand, *P. antipodarum* harbor extensive across-lake population genetic structure for both mitochondrial and nuclear genomes, but with significant mitonuclear discordance [21–24].

Nuclear genomes show strong clustering within lakes and differentiation among lakes, but similar mitochondrial haplotypes exist across multiple lakes [24]. This distribution of mitochondrial vs. nuclear variation allows for sampling asexual lineages that harbor closely related mitochondrial haplotypes in combination with the nuclear genomes from multiple lakes to estimate whether interactions between mitochondrial and nuclear genetic variation explain standing variation for metabolic rate and metabolic thermal plasticity among asexuals.

In the last 200 years, *P. antipodarum* has aggressively colonized aquatic ecosystems worldwide, to the extent that these snails have been identified as one of the “one-hundred worst” invasive taxa [25]. Like other invasive taxa, invasive *P. antipodarum* populations harbor low genetic diversity relative to the native populations [25, 26]. The widespread invasion, establishment, and successful monopolization of resources in the introduced aquatic habitat might be linked to adaptive phenotypic plasticity, among other factors, in *P. antipodarum* [25–28], but the underlying mechanisms remain unclear.

We quantified the mitonuclear contribution to variation in metabolic rate and thermal metabolic plasticity in native *P. antipodarum* and compared this variation to invasive lineages of *P. antipodarum*. Our findings demonstrate how temperature variation can reveal mitonuclear variation for metabolic rate within a species and suggest that a subset of this variation in metabolic plasticity may be favored in invasive lineages. We discuss our findings in the context of two competing hypotheses (increased intake hypothesis vs. compensation hypothesis) that have been proposed as explanations for associations between RMR, life-history traits and fitness and tested in a variety of invasive plant and animal taxa.

## MATERIALS AND METHODS

### Snail sampling, ploidy determination, and mtDNA assignment

In 2016 and 2018, snails were collected with kick nets from rocks and vegetation in the shallow regions of seven South Island New Zealand lakes (Alex, Brunner, Grasmere, Mapourika, Moeraki, Rotoroa, Selfe) that were known to harbor multiple mtDNA haplotypes amongst asexual *P. antipodarum* and have distinct across-lake nuclear genetic structure [22–24]. Native-range *P. antipodarum* populations harbor both sexual (diploid males and females) and asexual polyploid females, while invasive *P. antipodarum* are exclusively asexual [29–31]. In order to avoid conflating potential effects of ploidy and reproductive mode, we screened for and selected only triploid snails, which is the most common ploidy level for asexual *P. antipodarum* in both native and invasive populations [23, 31]. Flow cytometry using the Beckton Dickon FACS Aria in the Flow Cytometry Facility in the Carver College of Medicine at the University of Iowa was conducted following protocols in [29] using a single juvenile snail produced by a single female snail isolated from each field collected lineage to assign lineage ploidy, and from which we inferred reproductive mode [21]. Lineage in our study refers to the clonal descendants of these field-collected triploid, asexual snails.

To contrast these native asexual lineages with invasive lineages, we acquired triploid asexual North American invasive lineages from Center County (PA), Columbia River (WA), Muscureteang River (NJ), Snake River (ID), Broadman River (MI), the widely distributed US1 lineage, and a lineage from the “GeraardsB” European population [Geraardsbergen (Belgium)] detailed in Donne *et al*. (2020) [27]. A total of 90 triploid lineages were isolated from seven New Zealand lakes, and seven triploid lineages were isolated from the six US collection sites and the Belgian collection site.

The CHAOS DNA extraction protocol was used to extract genomic DNA from one individual in each of native and invasive lineages [32]. We then followed published procedures to sequence and assign mtDNA *cytochrome b* (*cytB*) haplotypes in this species [23, 24, 27]. We used the *cytB* primers and PCR/sequencing protocols described in Neiman *et al*. (2011) to sequence a 718 bp fragment of this gene from one individual from each of the lineages included in the study [23]. We manually checked the sequencing data (including chromatograms), trimmed the sequence to the 718 bp corresponding to the targeted region, and then used NCBI’s blastn “megablast” option to blast the 718 bp sequence that resulted to NCBI’s “nucleotide” database. Each of the new sequences matched at least one already characterized *cytB* haplotype at 99%+ identity. We assigned the new sequence to the haplotype that either matched the sequence at 100% identity or was the highest match if there was not 100% identity.

### Rationale for lineage selection

*P. antipodarum* populations (i.e., the community of potentially interbreeding individuals at a given locality) have marked among-lake nuclear genetic differentiation in New Zealand [24], and previous mtDNA-based studies of the phylogeography of *P. antipodarum* indicate that major genetic differences occur on the north/south axis of New Zealand [22, 23, 24]. We therefore used lake of origin as a proxy for nuclear genotype for native lineages. From the 90 asexual lineages that we sampled, we selected a subset of asexual lineages from three South Island New Zealand lakes—Lake Brunner, Lake Gunn and Lake Mapourika—because our sampled lineages from each of these lakes included each of three *cytB* mitochondrial haplotypes (1A, 20A and 37A). This allowed for a design with three mtDNAs each combined with two or three lake nuclear backgrounds for a total of eight native asexual lineages to be phenotyped for metabolic rate thermal plasticity. The invasive populations included in our study harbored either the 22A (GeraardsB, Muscureteang River, Broadman River, US1), 37A (Center County, Snake River) or 37D cytochrome b (*cytB*) mitochondrial haplotype (Columbia River).

### Experimental design

Snails were shipped on ice by overnight delivery from the University of Iowa. Upon arrival at the University of Nebraska-Lincoln, snails were housed in 0.5 L plastic cups at 16 °C on a 18 hr light/6 hr dark schedule, and fed *Spirulina* algae 3X per week for 10 days [19, 32]. Snails from each of the eight native lineages and seven invasive lineages described above were randomly assigned to one of two water temperature treatment groups, and each group was exposed to the temperature treatment for two weeks. Previous work on mollusks has shown that a two-week acclimation is sufficient to alter cellular responses to temperature [33, 34]. Our temperature treatments were chosen to represent a reasonable water temperature at the time of collection (16 °C; “control” temperature) in New Zealand, and a + 6 °C increase predicted for the year 2100 by an Intergovernmental Panel on Climate Change scenario (22 °C; stressful temperature) [35]. Invasive *P. antipodarum* have been shown to tolerate temperatures from 0 to 28 °C [26, 36, 37]. Thus, the temperatures used in our study are within the ecologically relevant range for this species. Water was changed 3X weekly. Mortality was checked daily, and animals that did not respond to a mechanical stimulus were recorded as dead and removed.

### Snail respirometry

Routine metabolic rate (RMR) was measured as the rate of oxygen consumption 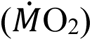 of snails at their respective acclimation temperature using the Oxygraph Plus System (Hansatech Instruments, Norfolk, UK) in 3 mL water-jacketed glass chambers equipped with a magnetic stirrer and Clark-type oxygen electrodes. The temperature of the respiration chambers was kept constant at either 16 °C or 22 °C using a Fisher Isotemp 4100 R20 refrigerated water circulator (Fisher Scientific, Hampton, NH). A two-point calibration of electrodes using air-saturated distilled water and sodium sulfite was used to establish 100% and 0% oxygen levels in the chamber, respectively. Ten biological replicates (i.e., 10 clonal snails) from each lineage and temperature combination were randomized across chambers and respirometry runs. Data were acquired and respiration rates were corrected for electrode drift using the OxyTrace+ software. After each approximately 2-hour individual respirometry run, snails were dissected to determine wet tissue mass, and a scaling co-efficient of 0.75 was used to calculate a mass-corrected RMR as RMR/mass^0.75^ [38].

### Statistical Analyses

We used the statistical package R version 2.15.1 for all statistical analyses (R Development Core Team 2011) [39]. For native lineages, we tested for the effects of temperature, mtDNA, and lake nuclear genetic variation on metabolism by applying a generalized linear model with temperature, lake nuclear genotype, mitochondrial haplotype, and their interactions as fixed factors and mass-corrected RMR as the response variable. For invasive lineages, we used two-way ANOVA to test for effects of lineage and temperature on mass-corrected RMR. Post hoc comparisons to test differences between group means were done using Tukey HSD tests that were corrected for multiple testing. We also contrasted mass-corrected RMR between invasive and native lineages under thermal stress using a nested ANOVA with temperature and status (native vs. invasive) as fixed factors and lineage as a nested factor within status. Unless otherwise indicated, data are shown as means ± standard errors of means (SEM).

## DATA AVAILABILITY

Supplemental files including all phenotype data are available at FigShare. Supplemental material available at Figshare: 10.6084/m9.figshare.23969031

## RESULTS

### RMR of Native Snails

Temperature, mitochondrial haplotype, and lake nuclear genotype had a significant three-way interaction effect on mass-corrected RMR of native asexual snails (*P* = 0.01) (Fig 1A, Fig 1C-E, Supplemental Fig1, Supplemental Fig 3A and 3B, Table 2, Supplemental Table 1). For snails from lakes Brunner and Mapourika, elevated temperature increased RMR (*P* = 4.47e^-06^ and *P* = 6.65e^-05^, respectively) (Table 2), but the magnitude of increase was dependent on mitochondrial haplotype (mito X temp, *P* = 0.03 and *P* = 0.03, respectively) (Fig 1A, Fig 1C, Fig 1E and Supplemental Fig 3A and 3B, Table 2). Snails from Lake Brunner with mitochondrial haplotype 37A and from Lake Mapourika with haplotype 20A maintained a similar RMR despite warming with no significant effect of temperature on RMR (*P* = 0.96 and *P* = 0.44, respectively.) On the other hand, all snails from Lake Gunn increased their RMR under warming (temp, *P* = 1.11e^-07^) with no effect of mitochondrial haplotype (mito, *P* = 0.42) and no interaction between temperature and mitochondrial haplotype (temp X mito, *P* = 0.99) (Fig 1A and 1D and Supplemental Fig 3A and 3B, Table 2). Thus, we find significant mitonuclear genetic variation for metabolic rate and metabolic rate plasticity among native asexual lineages of *P. antipodarum*, with some lineages recovering a RMR similar to their 16 °C RMR after a two-week acclimation to 22 °C.

**Figure 1:**
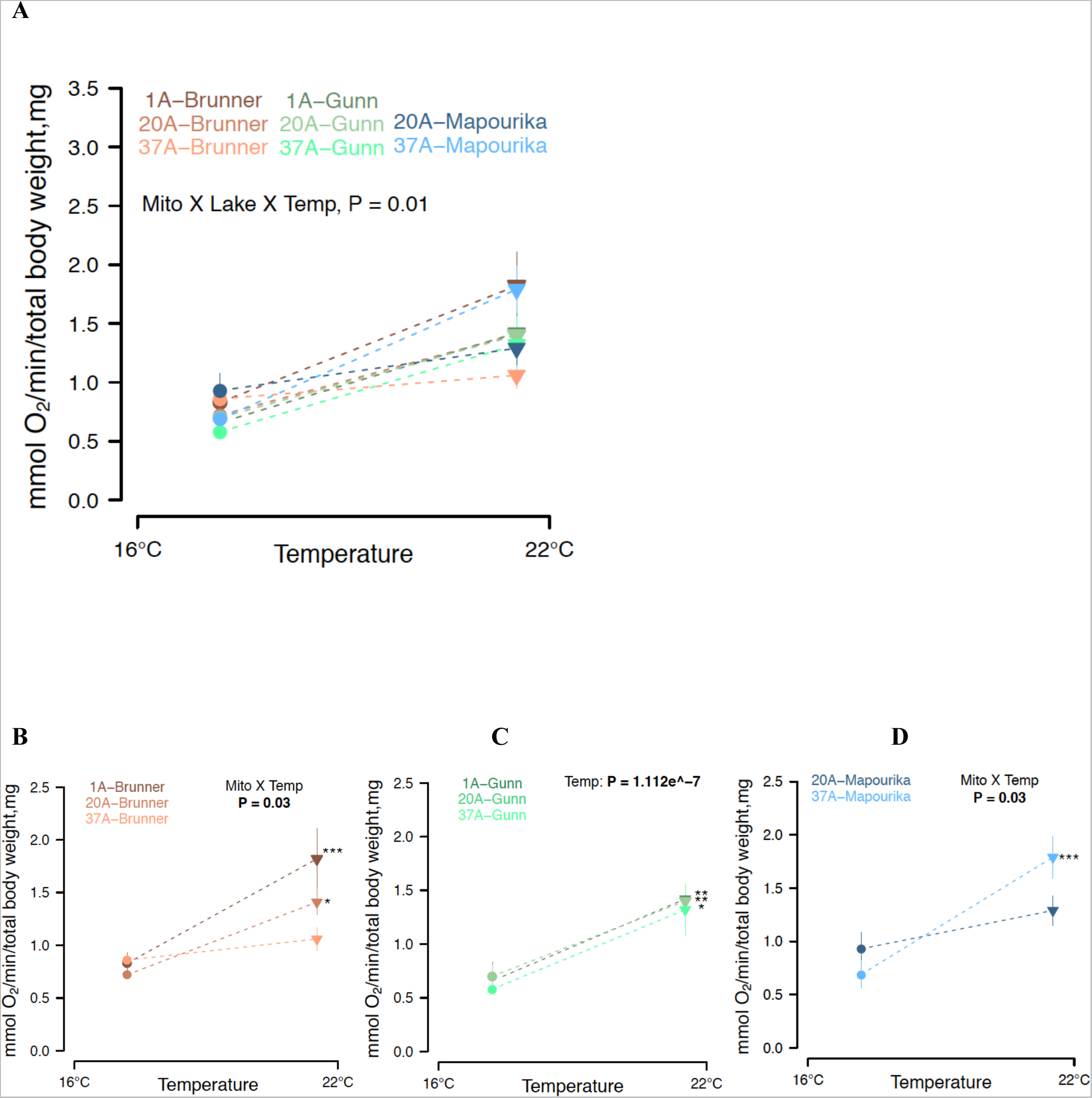
Routine Metabolic Rate (RMR) of native *Potamopyrgus antipodarum* exposed to different temperatures. RMR was measured in snail groups at their respective acclimation temperatures; i.e. 16 °C (closed circles) vs. 22 °C (closed triangles). Within each graph, asterisks indicate means are significantly different between temperature treatments (*p* < 0.05). Vertical bars represent SEM. N = 8–10 snails. C-E: RMR of native *P. antipodarum* lineages from lakes Brunner, Gunn and Mapourika with mitochondrial haplotypes 1A, 20A and 37A

### RMR of Invasive Snails

Lineage had a significant effect on mass-corrected RMR (lineage, *P* = 0.02) in a 2-way ANOVA model with no significant effect of temperature or its interaction with lineage (temp, *P* = 0.52, temp X lineage, *P* = 0.75) (Fig 2, Supplemental Fig 1A, Supplemental Fig 2A and 2B, Table 2). At 16 °C, there was no significant difference in RMR among the lineages (lineage, *P* = 0.53), but at 22 °C snails from Snake River had significantly lower RMR than snails from Broadman River (*P* = 0.0007) and US1 snails (*P* = 0.005) (Fig 2, Supplemental Fig 2A and 2B, Supplemental Table 1). One-way ANOVA results for the effect of temperature in each individual lineage were non-significant (all *P*> 0.05), demonstrating that all the lineages were non-plastic for RMR in response to temperature with all lineages recovering a RMR similar to their 16 °C RMR after a two-week acclimation to 22 °C (Table 2).

**Figure 2:**
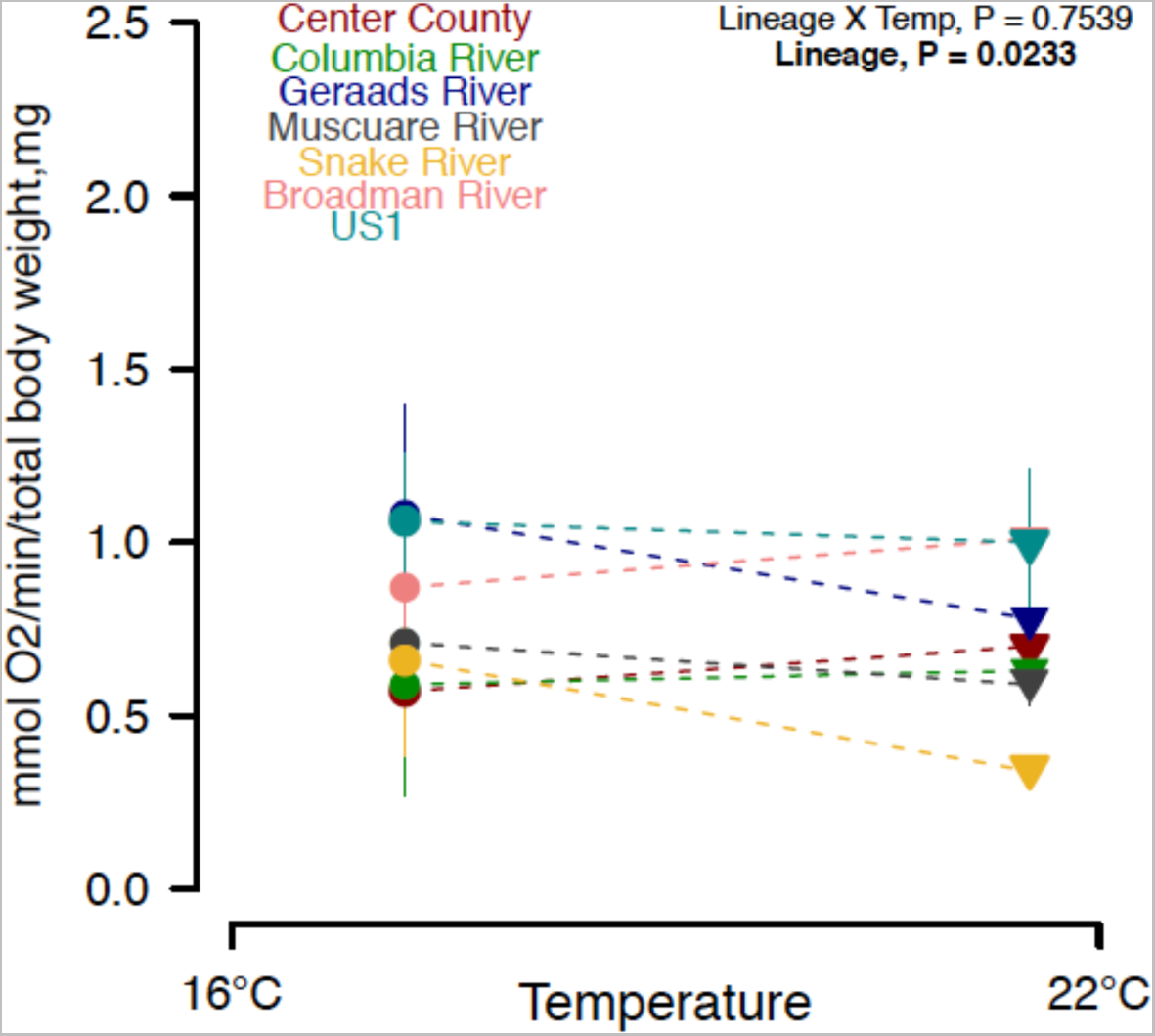
Routine Metabolic Rate (RMR) of invasive *Potamopyrgus antipodarum* exposed to different temperatures. RMR was measured in snail groups at their respective acclimation temperatures; i.e. 16 °C (closed circles) vs. 22 °C (closed triangles). Vertical bars represent SEM. N = 8–10 snails.

**Figure 3:**
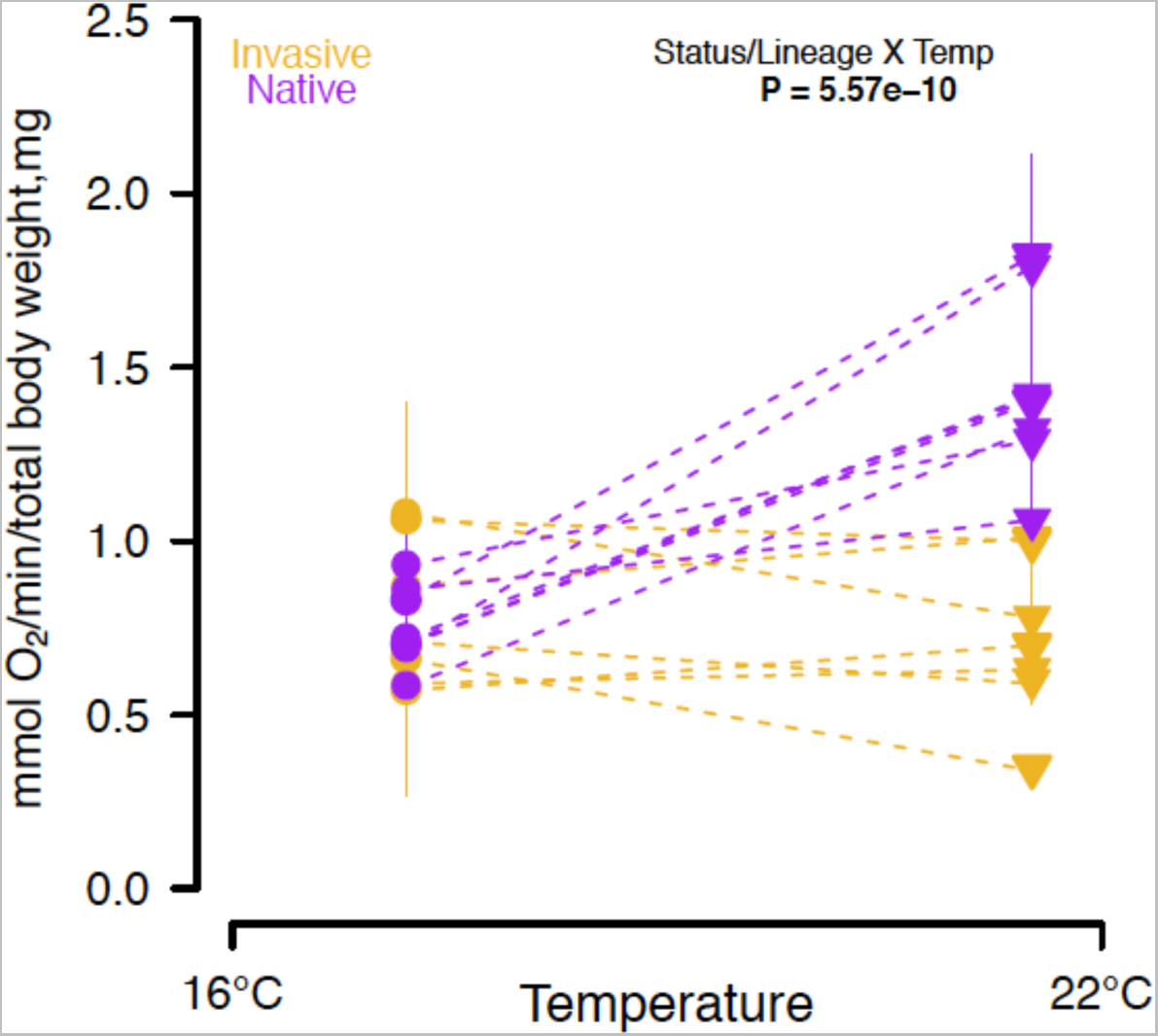
Comparision of Routine Metabolic Rate (RMR) of invasive (golden yellow) and native (purple) *P. antipodarum* exposed to different temperatures. RMR were measured in snail groups at their respective acclimation temperatures; i.e. 16 °C (closed circles) vs. 22 °C (closed triangles). Asterisks indicate means are significantly different between temperature treatments (*p* < 0.05). SEM. N = 8–10 snails.

### Comparison of RMR between Native and Invasive Snails

Combining data from both native and invasive snails, we found a significant effect of native vs. invasive status (*P* = 0.0016), temperature (*P* = 4.46 e^-09^), and their interaction (status X temp, *P* = 5.57e^-10^) on mass-corrected RMR (Fig 3, Supplemental Fig 1A and 1B, Table 1). Lineage as a nested factor within status had a significant effect on RMR at 22 °C (Lineage/Status, *P* = 0.0002) but not at 16 °C (Lineage/Status, *P* = 0.5672) (Table 1). At 16 °C, the mean RMR of native snails was not significantly different from the mean RMR of invasive snails (*P* = 0.81). By contrast, at 22 °C the mean RMR of native snails was significantly different and nearly twice that of the invasive snails (*P* = 2e^-16^) (Table 1).

**Table 1:**
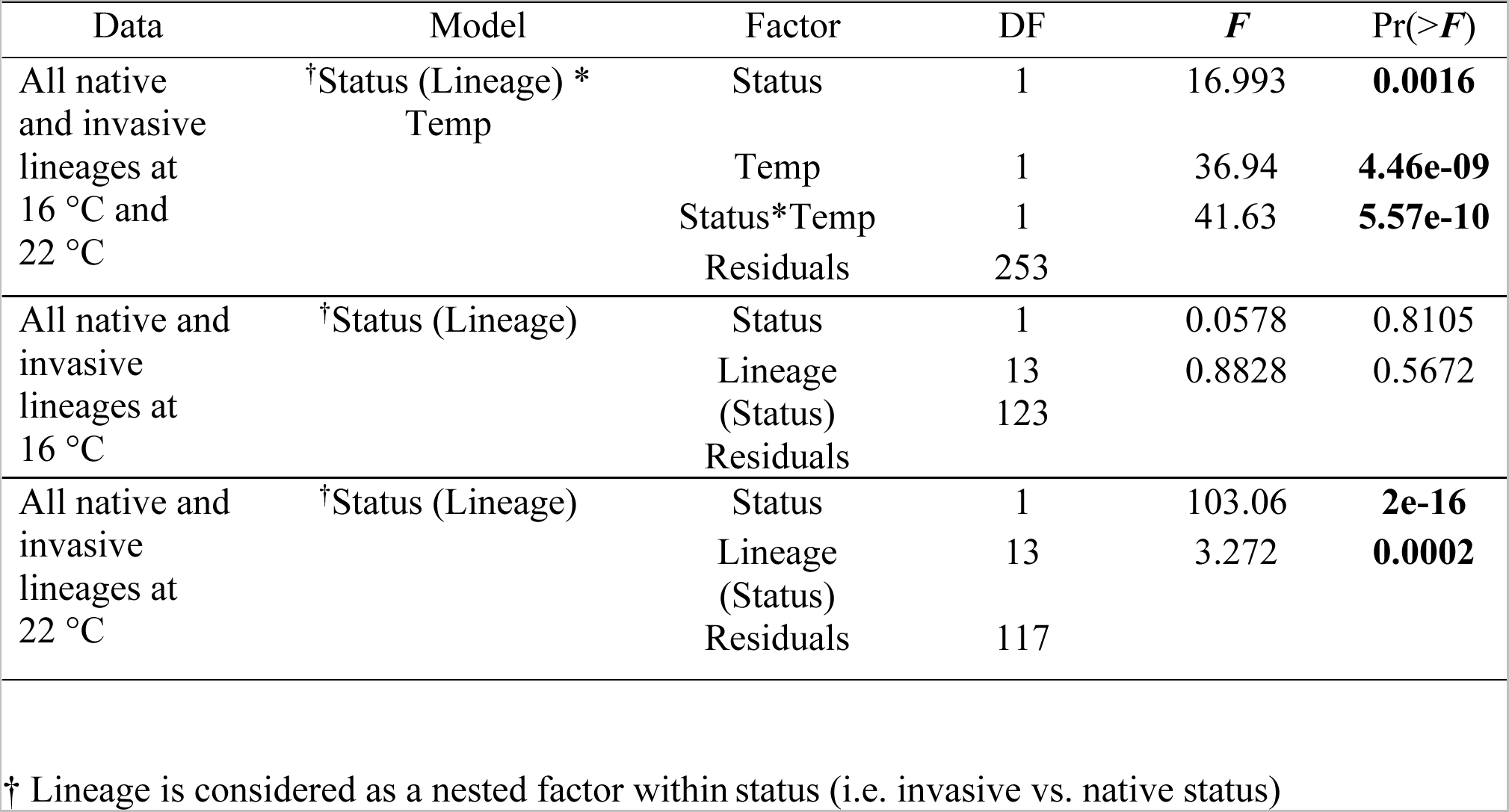
Mixed-model ANOVA results for the effects of exposure temperature (16 °C vs. 22 °C), status (invasive vs, native), and the interaction of these factors on mass-corrected routine metabolic rate (RMR) in *Potamopyrgus antipodarum*. Lineage was considered as a random nested factor within status. Numbers in bold indicate *p* < 0.05).

**Table 2:** ANOVA results for the effects of exposure temperatures (16 °C vs. 22 °C), lake of origin, mitochondrial haplotype, and the interaction of these factors on mass-corrected routine metabolic rate (RMR) in native *Potamopyrgus antipodarum*. For invasive *P. antipodarum*, exposure temperature and lineage were used as factors.

| Lineage | Model | Factor | DF | *F* | Pr(>*F*) |
| --- | --- | --- | --- | --- | --- |
| Native Lineages |  |  |  |  |  |
| All | Lake*Mito* Temp | Lake | 2 | 1.2032 | 0.3035 |
|  |  | Mito | 2 | 2.5608 | 0.08108 |
|  |  | Temp | 1 | 84.3758 | 7.49E-16 |
|  |  | Lake: Mito | 3 | 1.1018 | 0.35087 |
|  |  | Lake: Temp | 2 | 0.2928 | 0.74667 |
|  |  | Mito: Temp | 3 | 1.2732 | 0.28335 |
|  |  | Lake:Mito:Temp | 2 | 3.575 | **0.01581** |
|  |  | Residuals | 132 |  |  |
| Brunner | Mito*Temp | Mito | 2 | 3.2331 | **0.04764** |
|  |  | Temp | 1 | 26.359 | **4.47E-06** |
|  |  | Mito: Temp | 2 | 3.6421 | **0.03323** |
|  |  | Residuals | 51 |  |  |
| Gunn | Mito*Temp | Mito | 2 | 0.8656 | 0.4274 |
|  |  | Temp | 1 | 39.1032 | **1.11E-07** |
|  |  | Mito: Temp | 2 | 0.0075 | 0.9925 |
|  |  | Residuals | 47.8 |  |  |
| Mapourika | Mito*Temp | Mito | 1 | 0.8034 | 0.37638 |
|  |  | Temp | 1 | 20.7103 | **6.52E-05** |
|  |  | Mito: Temp | 1 | 4.8897 | **0.03384** |
|  |  | Residuals | 34 |  |  |
| Invasive Lineages |  |  |  |  |  |
| All | Lineage* Temp | Lineage | 6 | 2.5601 | **0.02334** |
|  |  | Temp | 1 | 0.4103 | 0.52317 |
|  |  | Lineage:Temp | 6 | 0.5693 | 0.75398 |
|  |  | Residuals | 109 |  |  |
| Snake River | Temp | Temp<br>Residuals | 1<br>16 | 1.5894 | 0.2255 |
| Columbia River | Temp | Temp<br>Residuals | 1<br>17 | 0.0176 | 0.8961 |
| Center County | Temp | Temp<br>Residuals | 1<br>17 | 0.716 | 0.4092 |
| Muscuare River | Temp | Temp<br>Residuals | 1<br>18 | 0.2947 | 0.5939 |
| Geraads River | Temp | Temp<br>Residuals | 1<br>13 | 1.0214 | 0.3306 |
| US1 | Temp | Temp<br>Residuals | 1<br>12 | 0.035 | 0.8548 |
| Broadman River | Temp | Temp<br>Residuals | 1<br>16 | 0.4808 | 0.498 |

## DISCUSSION

We found that nuclear genetic variation (via lake population as a proxy), mitochondrial genome, and their intergenomic mitonuclear interaction significantly shape the thermal plasticity of metabolic rate in asexual *P. antipodarum*. Like other organismal traits, RMR has a genetically complex architecture with estimates of heritability ranging from 0-0.72 [40 and references therein]. Coupled transmission of the nuclear and mitochondrial genomes without recombination in asexual *P. antipodarum* lineages should reduce the efficacy of natural selection in both nuclear [41] and mitochondrial genomes [17]. Yet, the permanent linkage between the nuclear and mitochondrial genomes in these lineages could also facilitate mitonuclear coevolution via clonal selection for compatible combinations of mitochondrial and nuclear variants that maintain or increase fitness in asexual lineages [19, 42, 43]. This possibility is of interest in light of the results of Greimann *et al*. (2020), who reported no difference in mitochondrial function and RMR between sexual vs. asexual lineages of native *P. antipodarum* [19] despite evidence for the accumulation of putatively harmful mutations in the mitochondrial genomes of asexual *P. antipodarum* [44]. It is possible that clonal selection favors the persistence of asexual lineages with nuclear variants that compensate for these mtDNA mutations and maintain RMR [19, 44]. Consistent with effective natural selection, we found no significant genetic variation in metabolic rate among the native asexual lineages at 16 °C, a common temperature experienced by these snails in their native range. By contrast, chronic exposure and acclimation to an elevated temperature of 22 °C revealed significant among-lineage genetic variation in RMR in two of the lakes. This hidden or “cryptic” genetic variation that is revealed to natural selection under thermal stress might be an important source of variation that could allow *P. antipodarum* populations to respond to environmental change [45].

Acute exposure to temperature increases metabolic rates of ectotherms but, over time, physiological change via acclimation or because of thermal stress can counter this thermodynamic effect [8]. Out of the eight native asexual lineages used in this study, six lineages had a significantly elevated RMR after two weeks at elevated temperature (i.e., they had thermally plastic RMR.) In contrast, a subset of lineages from Lake Brunner and Lake Mapourika showed complete thermal compensation of RMR, such that the acclimated RMR at 22 °C was not different from their respective RMR at 16 °C (i.e., RMR was not thermally plastic). Lineages with the same *cytB* mitochondrial haplotype showed varying degrees of thermal plasticity of RMR that depended on lake nuclear genome, which manifested as a significant mito-nuclear-temperature or GxGxE interaction. Q_10_ (or ‘acclimated’ Q_10_ in our case) is a measure of temperature sensitivity of a physiological trait [8]. Mitotype 37A, when combined with the nuclear background of Lake Mapourika, was most plastic for RMR (Q_10_ = 4.82), while the RMR of haplotype 37A with the nuclear background of Lake Brunner was non-plastic (Q_10_ = 1.43) (Supplemental Table 2). This result underscores the importance of mitonuclear genetic interactions as a source of variation in metabolic plasticity, especially under temperature scenarios expected under climate change.

At this time, we still lack a general rule or predictive framework for the relationship between metabolic rate, metabolic plasticity, and fitness. To gain insight and provide a context for our observation of mitonuclear variation for metabolic thermal plasticity in native asexual snails, we tested whether invasive asexual lineages represented a subset of this phenotypic variation that has been successful in invading globally. In contrast to the native lineages, no invasive asexual lineages of *P. antipodarum* were plastic for RMR in response to elevated temperature—a response similar to a small subset (two of eight) of native asexual lineages. At 16 °C, the RMR of invasive and native snails were indistinguishable, while the RMR of most native lineages was about twice that of invasive lineages at 22 °C. This result suggests that under elevated temperatures, invasive asexual lineages did not or were not able to allocate additional energy for maintenance metabolism, resulting in what appeared as complete thermal compensation (Q_10_ ∼ 1) at the end of the 2-week acclimation period.

*P. antipodarum* is an aggressive global invader of aquatic ecosystems worldwide and is characterized by very low genetic diversity in its invaded range [25–28]. Only two mitochondrial haplotypes have been identified in Europe, whereas more than 50 mitochondrial haplotypes have been characterized in the native New Zealand lakes [24, 28, 46, 47]. Nuclear genetic diversity is somewhat higher than mitochondrial diversity in the invasive range in Europe [28, 46], but does not come close to the high nuclear genetic variation amongst asexual *P. antipodarum* in New Zealand [24, 48]. The genetic diversity observed in Europe is low enough to suggest post-invasion diversification [26, 47] rather than the multiple separate origins of clones that the New Zealand data indicate [24, 48]. Outside of the diversification linked to mutation post-invasion, invasive lineages of *P. antipodarum* seem to represent a subset of native variation that is beneficial in the setting of invasion [27]. Indeed, both North American and European invasive lineages harbor mtDNA haplotypes common on the North Island of New Zealand [23, 47]. The nuclear genotypes of the European invaders in particular harbor genotypes similar to those found in both of New Zealand’s major islands, while North American invaders seem more similar to the North Island [24]. Altogether, an even more thorough and comprehensive survey than that provided in Donne *et al*. (2020) will ultimately be needed to completely reconstruct the genetic relationships between and across native and invasive populations of *P. antipodarum*.

Nevertheless, our observations indicate that the absence of metabolic thermal plasticity in invasive *P. antipodarum* represents a subset of the phenotypic and potentially mitonuclear genetic variation for metabolic rate that exists in the native range.

There are two potential explanations invoked to explain the well-documented relationships between metabolic rate, pace of life history, and invasion [1]. The “increased intake hypothesis” posits that individuals with higher maintenance costs also have more metabolic machinery and higher capacity to enhance fitness-related functions such as aerobic performance, mobility, and thermogenesis [1, 49, 50]. By contrast, the “compensation hypothesis” posits that individuals with higher maintenance costs have less energy surplus to allocate to other physiological functions, resulting in trade-offs with growth, reproduction, and immune defense [1, 50, 51]. Our metabolic rate data from invasive *P. antipodarum* support the compensation hypothesis, especially when taken together with previous studies of life-history trait variation in *P. antipodarum* [27]. The ability of invasive snails to maintain a low metabolic rate at higher temperature means that out of a finite organismal energy budget, a lower proportion of metabolic energy is allocated for routine or maintenance metabolism at higher temperatures in invasive snails. This in turn makes investment of a larger proportion of energy possible for other life-history traits, which is intriguing in light of the documented earlier reproductive maturity and lower growth rates but nevertheless similar adult size of invasive snails when compared to native lineages [27]. The metabolic rate data in our study offers a possible link for observed differences in reproductive strategies of invasive and noninvasive lineages. Together, these results point to variation in life-history trade-offs between invasive and non-invasive lineages of *P. antipodarum*. This pattern of flexibility in life-history traits – which could facilitate colonization - is also observed in other invasive species [52, 53].

Metabolic rate is an emergent property resulting from interactions among various physiological functions within an individual organism. These organism-level functional interactions are expected to scale up and shape higher-level ecological processes and patterns. For example, Shuster *et al*. (2021) investigated key theoretical predictions about relationships between metabolism and demography in a marine invertebrate (*Bugula neritina*) and found that populations with relatively low metabolic rates had relatively high carrying capacities [54]. Thus, it is possible that the ability of invasive *P. antipodarum* to maintain low metabolic rates under environments that are expected to increase maintenance costs and metabolic rates could help explain their ability reach population densities up to 800,000 individuals/m^2^ in the invasive range [55, 56] compared to 10,000-14,000 individuals/m^2^ in its native range in New Zealand [57].

Lower metabolic rates may be selected for in environments where reduced energetic requirements are advantageous such as empty habitats, low predation pressure, or temperature stress [58]. Invasive *P. antipodarum* are very successful with respect to the former: they typically invade during the early stages of ecological succession in an aquatic ecosystem [30]. *P. antipodarum* also has few predators and no known parasites in the invasive range, meaning that reproduction may be the main fitness determinant for invasive populations [30, 59]. Taken together with these studies, our data suggest a potential connection between relatively low allocation of energy to maintenance metabolism while maximizing lifetime reproductive output and successful colonization of non-native habitats by invasive *P. antipodarum*.

We also observed that RMR differed among invasive lineages of *P. antipodarum* but only under thermal stress and driven primarily by the low of RMR of invasive *P. antipodarum* from Snake River (ID) relative to other lineages. Evidence for variation among other life-history traits in invasive *P. antipodarum* populations has been previously reported [27, 60]. In particular, invasive snails collected from Snake River (ID) grew more slowly than other invasive populations in a common-garden experiment at 16 °C [27] but had higher growth rates at 24 °C [60], an elevated environmental temperature similar to where we observed the lowest RMR in snails from this same location. Altogether, these results suggest that the Snake River population is a “Master-of-Some” situation [27] where their ability to maintain a low metabolic rate at high temperatures enables plasticity for other life-history traits that may increase their fitness.

It is important to note the potential limitations of our study design regarding our use of only two temperature treatments. In our study, exposure to 22 °C for two weeks likely represents a stressful temperature for native lineages. Chronic temperature exposure higher than 22 °C results in mass mortality in native *P.antipodarum* lineages (personal obs.) However, numerous studies have reported that species in their invasive range show eurythermality, i.e., the ability to maintain physiological function across an extensive range of temperatures [61, and references therein]. In natural and laboratory settings, a wide temperature tolerance has also been documented for *P. antipodarum* ranging from 2 °C [62] to 34 °C [63]. Thermal performance curves (7 °C - 35 °C) for RMR for invasive *P. antipodarum* collected from the western US demonstrate that acute exposure to elevated temperature does increase oxygen consumption with max RMR at 28 °C (T_opt_), beyond which RMR declined sharply [64]. We observed that after two weeks at elevated temperature, the Q_10_ is ∼ 1 (i.e., the thermal reaction norm is flat or non-plastic) for invasive *P. antipodarum*, indicating significant acclimation in invasive *P. antipodarum* at 22 °C through thermal compensation. This outcome points to a nuanced yet critical point: achieving a flat reaction norm for RMR during thermal acclimation is only possible when underlying sub-organismal physiological, cellular, and biochemical processes are plastically responding to temperature. Schulte *et al*. 2011 elegantly describes this as leading to “the apparently absurd conclusion that the only way to achieve a reaction norm demonstrating a lack of plasticity is for the organism to exhibit substantial plasticity” [65]. Thus, it may be that extensive cellular and physiological plasticity may be underlying eurythermality and the thermal compensation of metabolic rates that we observed to support plastic life-history traits that have been implicated in the invasive success of *P. antipodarum* [25, 46].

## ACKNOWLEDGEMENTS

The flow cytometry data that we used to identify triploid lineages were obtained at the Flow Cytometry Facility, which is a Carver College of Medicine / Holden Comprehensive Cancer Center core research facility at the University of Iowa. The facility is funded through user fees and the generous financial support of the Carver College of Medicine, Holden Comprehensive Cancer Center, and Iowa City Veteran’s Administration Medical CentFIGURES.

