## Supplemental figures and Tables for "Mitochondrial-nuclear variation for metabolic plasticity with potential consequences for invasion success in New Zealand mud snails"

**Supplemental Figure 1**. Violin plots show a significant effect of acclimation temperature on mass-corrected routine metabolic rate (RMR) of (A) native but not (B) invasive lineages of *Potamopyrgus antipodarum*. RMR was measured in replicate groups of 3 snails at their respective acclimation temperatures of 16 °C or 22 °C. N= 8-10.

**A.** RMR in Native Lineages

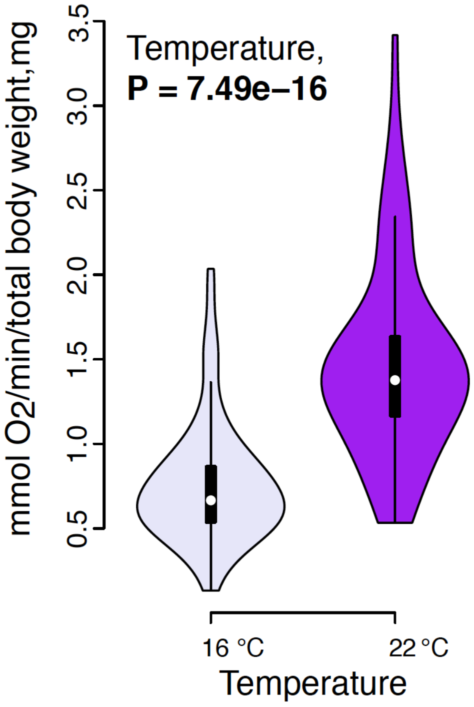


**B.** RMR in Invasive Lineages

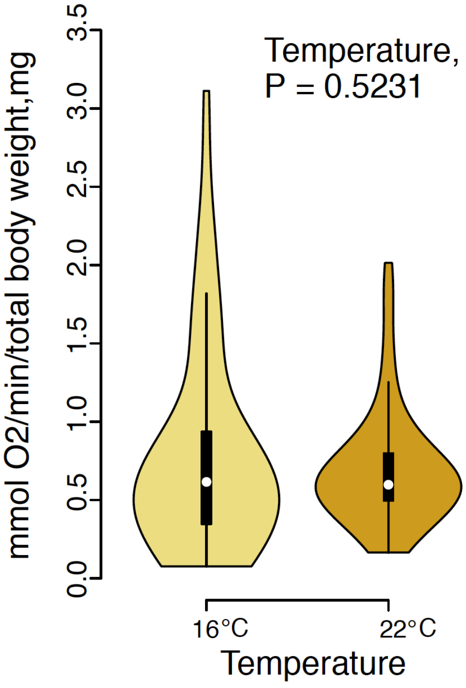

**Supplemental Figure 2.** Box plots show significant effects of an interaction between mitochondrial type and lake (a proxy for nuclear genotype) on mass-corrected routine metabolic rate (RMR) of native lineages of *P. antipodarum* at 22 °C (B) but not at 16 °C (A). RMR was measured in 8-10 replicate groups of three snails per lineage at their respective acclimation temperatures.

**A.**

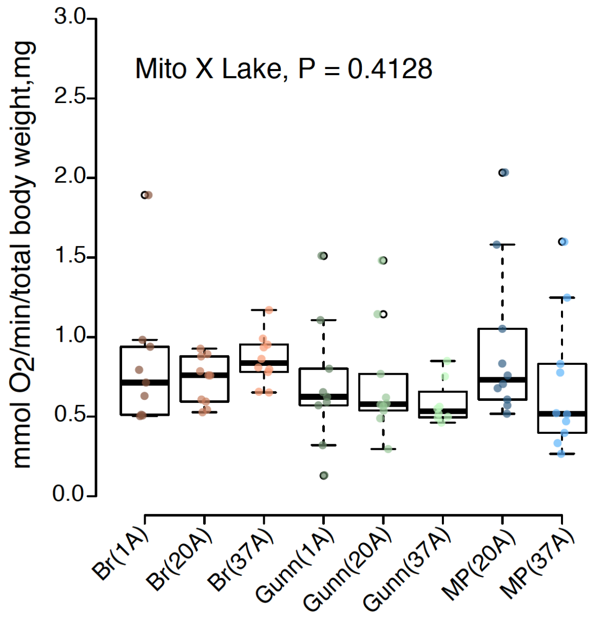


**B.**

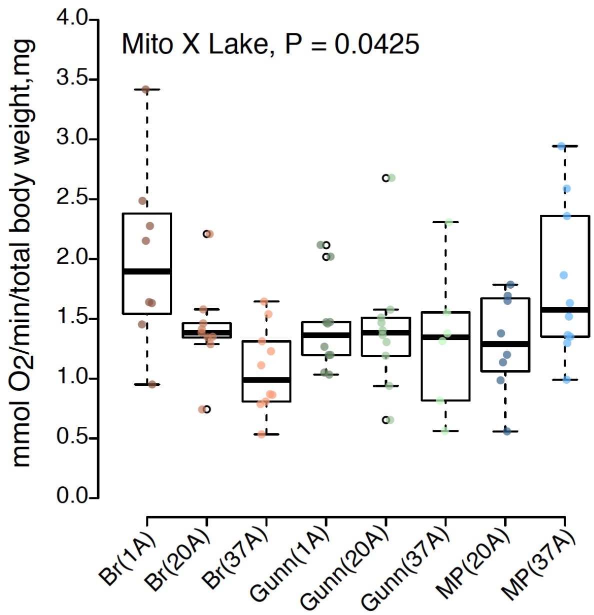

**Supplemental Figure 3.** Box plots show significant effects of genetic lineage on mass-corrected routine metabolic rate (RMR) of invasive *P. antipodarum* at 22 °C (B) but not at 16 °C (A). RMR was measured in 8-10 replicate groups of three snails per lineage at their respective acclimation temperatures.

**A.**

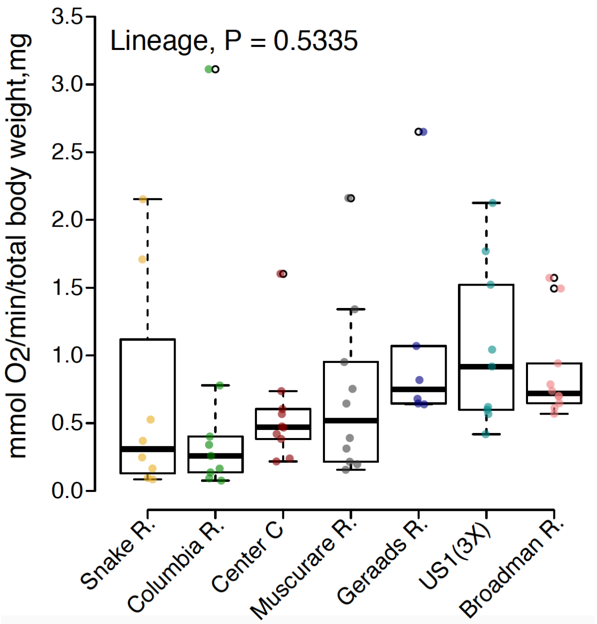


**B.**

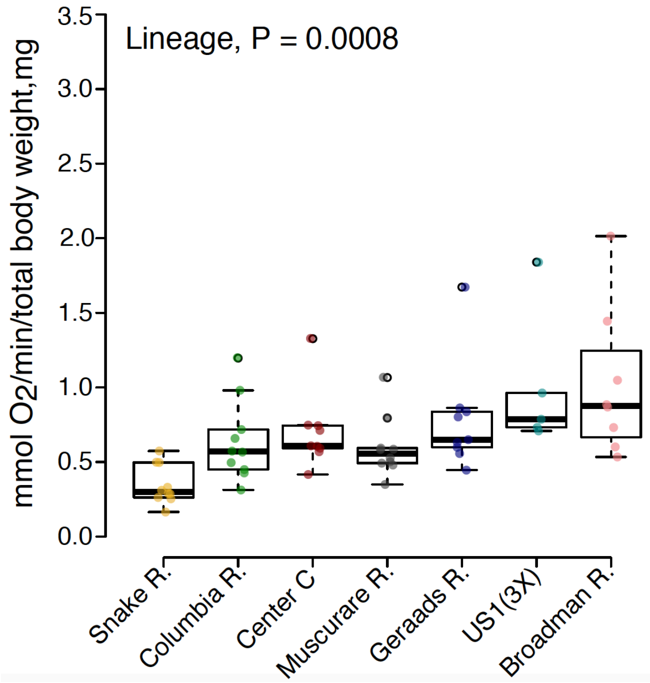

**Supplemental Table 1.** ANOVA for the effects of lake of origin (a proxy for nuclear genotype) and mitochondrial haplotype in native *P. antipodarum* and for the effect of lineage in invasive *P. antipodarum* on RMR of snails acclimated and measured at 16 °C and 22 °C. Significant *P* values < 0.05 are marked in bold.

| | Model | Factor | DF | F-value | Pr(>F) |
|---|---|---|---|---|---|
| **Native Lineages** | | | | | |
| At 16 °C | Lake*Mito | Lake | 2 | 1.5368 | 0.2224 |
| | | Mito | 2 | 0.2592 | 0.7724 |
| | | Lake: Mito | 3 | 0.9685 | 0.4128 |
| | | Residuals | 68 | | |
| At 22 °C | Lake*Mito | Lake | 2 | 1.5206 | 0.2263 |
| | | Mito | 2 | 1.1974 | 0.3086 |
| | | Lake: Mito | 3 | 2.8820 | **0.0426** |
| | | Residuals | 64 | | |
| **Invasive Lineages** | | | | | |
| At 16 °C | Lineage | Lineage | 6 | 0.8552 | 0.5335 |
| | | Residuals | 55 | | |
| At 22 °C | Lineage | Lineage | 6 | 4.5713 | **0.0008** |
| | | Residuals | 54 | | |

**Supplemental Table 2.** Mean RMR and calculated temperature quotient, $Q_{10}$ , for mass-corrected routine metabolic rate (RMR) in native and invasive *P. antipodarum* acclimated and measured at 16 °C and 22 °C.

| Type/ Lake | Lineage | Temperature, °C | Routine metabolic rate (RMR), (nmol/min/g wet tissue) | Temperature coefficient, $\mathbf{Q_{10}}$ $(\frac{\mathbf{R2}}{\mathbf{R1}})^{\frac{\mathbf{10}}{\mathbf{(T2-T1)}}}$ |
|---|---|---|---|---|
| **Native Lineages** | | | | |
| Brunner | Brunner -20A | 16 | 0.727 | 3.04 |
| | | 22 | 1.418 | |
| | Brunner-37A | 16 | 0.860 | 1.43 |
| | | 22 | 1.069 | |
| Gunn | Gunn-1A | 16 | 0.701 | 3.27 |
| | | 22 | 1.428 | |
| | Gunn-20A | 16 | 0.706 | 3.16 |
| | | 22 | 1.408 | |
| | Gunn-37A | 16 | 0.585 | 3.88 |
| | | 22 | 1.32 | |
| Mapourika | Mapourika-20A | 16 | 0.934 | 1.73 |
| | | 22 | 1.297 | |
| | Mapourika-37A | 16 | 0.696 | 4.82 |
| | | 22 | 1.824 | |
| **Invasive Lineages** | | | | |
| | Snake River | 16 | 0.669 | 0.33 |
| | | 22 | 0.346 | |
| | Columbia River | 16 | 0.595 | 1.12 |
| | | 22 | 0.638 | |
| | Center County | 16 | 0.571 | 1.41 |
| | | 22 | 0.702 | |
| | Muscuare River | 16 | 0.712 | 0.74 |
| | | 22 | 0.597 | |
| | Geraads River | 16 | 1.083 | 0.58 |
| | | 22 | 0.784 | |
| | US1(3X) | 16 | 1.064 | 0.90 |

| | | | |
|---|---|---|---|
| | 22 | 1.005 | |
| Broadman River | 16 | 0.876 | 1.27 |
| | 22 | 1.015 | |
